# From Chewing to Chirping: The Misophonia Audiovisual Trigger Archive (MATA)

**DOI:** 10.1101/2025.09.09.675007

**Authors:** Sewon Oh, Katherine Palmer, Danielle Sabatina, Alina Pietrini, Christian O’Reilly, Svetlana V. Shinkareva

## Abstract

Misophonia is an emerging condition in which everyday sounds, such as chewing, sniffing, or tapping, evoke disproportionately intense emotional and physiological responses. Despite growing recognition of its clinical significance, progress in understanding misophonia has been hindered by the limited availability of standardized and ecologically valid stimulus sets. Here, we present a large, open-access archive of 1,300 five-second audiovisual clips spanning 12 empirically validated categories of misophonic triggers. This resource extends beyond orofacial movement-related sounds to include a diverse array of real-world triggers, and its audiovisual format enables systematic investigation of how visual context shapes responses to misophonic sounds. The archive lowers the barrier for laboratories to study misophonia, promotes reproducibility across sites, and offers applications ranging from crowdsourced assessments of population-level sensitivities to machine learning approaches for automated trigger detection. By providing the largest and most diverse audiovisual misophonia stimulus repository to date, this resource is designed to accelerate mechanistic, clinical, and translational research on misophonia and related sensory-emotional phenomena.

## Background & Summary

### Context

Misophonia is an emerging condition in which everyday sounds, such as chewing, sniffing, or tapping, evoke disproportionately intense emotional and physiological responses [1, 2]. Far from being a minor annoyance, these sounds can elicit acute anger, distress, and autonomic arousal, often driving individuals to engage in avoidance or escape behaviors [e.g., 3]. Recent population-based estimates suggest that nearly one in twenty individuals in the United States experience misophonia [4]. Although oral and nasal sounds are most frequently reported, virtually any repetitive environmental noise can become a trigger. Misophonia has been linked to impaired quality of life [5], elevated anxiety in youth [6], and heightened risk of self-harm in young adults [7]. Together, these findings underscore misophonia as a prevalent and clinically significant condition warranting systematic scientific investigation.

### Motivation for Creating this Dataset

Most laboratory studies of misophonia have relied on orofacial movement-related sounds, such as chewing, to reliably elicit misophonic responses. While these sounds provide a strong experimental foundation, they represent only a subset of the diverse and idiosyncratic triggers reported by individuals with misophonia. A smaller number of investigations have incorporated person-specific triggers [e.g., 8, 9-11], underscoring the individualized nature of the condition. Using person-specific stimuli offers several advantages: it allows researchers to capture heterogeneity across individuals, provides a more sensitive test of mechanistic hypotheses, and reflects the ecological reality that responses to personally salient triggers are often more robust than to triggers that are more prevalent across individuals. More broadly, expanding research beyond orofacial movement-related sounds avoids over-reliance on a single stimulus category and enables a more comprehensive understanding of misophonia.

Despite these advantages, the use of individualized or ecologically diverse triggers has been limited by practical barriers. Generating stimuli is resource-intensive and time-consuming, constraining their broader adoption in experimental studies. Several important efforts have begun to address this gap. The Free Open-Access Misophonia Stimuli database [FOAMS; 12] provides a standardized set of 32 trigger sounds from eight categories, such as breathing and typing. The Sound-Swapped Video database [SSV; 13] introduced an innovative audiovisual design, pairing 18 misophonic trigger sounds with either their original video sources (e.g., chewing sounds paired with chewing videos) or with positively attributable visual sources (e.g., the same chewing sound paired with tearing paper). This approach has enabled direct tests of how visual context shapes misophonic responses. Yet, despite these advances, existing resources remain limited in scope, diversity, and do not capture the full range of audiovisual triggers encountered in daily life.

Here, we present a large, publicly available archive of 1,300 audiovisual stimuli spanning multiple categories of misophonic triggers. It addresses a critical bottleneck in the field: the limited availability of standardized and diverse trigger stimuli. By extending beyond orofacial movement-related sounds to include a broader array of real-world triggers (e.g., Styrofoam scratching, Velcro tearing), our archive provides researchers with a flexible tool for studying misophonia. The breadth of sound categories included ensures that the archive can capture at least some degree of individual difference in misophonic triggers, which is essential given the highly idiosyncratic nature of the condition. Critically, the audiovisual format offers an important advance over auditory-only collections: it allows for systematic investigation of how visual context shapes responses to trigger sounds, reflecting the inherently multisensory nature of misophonic experiences. This resource is designed to accelerate progress in uncovering the mechanisms and broader impact of misophonia.

### Previous usage

Using a subset of these stimuli, we examined both physiological and subjective responses to misophonia triggers [11]. Participants with and without misophonia were exposed to sounds, silent videos, and mental imagery of trigger, aversive, and non-aversive stimuli. Five physiological signals were recorded: facial muscle activity (two sites), skin conductance, heart rate, and temperature. Our findings demonstrate that misophonia triggers, relative to aversive stimuli, can be reliably distinguished through physiological responses. Moreover, misophonia and control groups could be differentiated across auditory, visual, and imagery conditions, underscoring the need for a comprehensive audiovisual stimulus repository.

### Potential Reuse Value

Beyond its immediate utility, this archive has several important implications for advancing misophonia research. First, it simplifies the process of locating or generating stimuli, thereby lowering the entry barrier for laboratories that may not have the resources to record and validate their own sets. Second, by offering a standardized and openly accessible resource, the archive promotes reproducibility across studies and supports the development of treatment protocols that can be tested and compared across sites. Finally, although designed for research, this resource could also be of interest to non-researchers, who may explore it for education, awareness, advocacy purposes, or for clinical applications.

The availability of a large-scale audiovisual archive also opens up novel applications. For example, the stimuli could be used in crowdsourced platforms, where individuals rate the degree of unpleasantness or social disruption associated with different triggers. Such collective data could enrich our understanding of trigger variability and population-level sensitivities. The archive also has potential utility in machine learning applications, such as training algorithms to automatically detect misophonic sounds, an approach that could inform assistive technologies (e.g., earphones that filter or attenuate identified triggers, as suggested in prior research). Because of its large size, it also enables for deep learning experiments, for example, synthesizing individualized triggers based on responses to known triggers or AI-assisted alteration of stimulus property inspired by the approach proposed with SSV. Moreover, the archive may prove useful in studying other sensory-emotional phenomena, offering a template for exploring how multisensory stimuli influence affective responses beyond misophonia.

## Methods

### Stimulus Categories

Stimuli were organized according to 12 broad categories derived from the Duke-Vanderbilt Misophonia Screening Questionnaire [DVMSQ; 14]. The DVMSQ includes a structured taxonomy of common misophonic triggers, grouping them into domains such as oral/nasal sounds (e.g., chewing, sniffing), environmental sounds (e.g., tapping, typing), and human movement-related sounds (e.g., foot shuffling, pen clicking). By aligning our archive organization with these empirically validated categories, we ensured that the stimulus set reflects the breadth of triggers most frequently endorsed in clinical and research samples, while also facilitating consistency and comparability with prior and future work using the DVMSQ framework.

### Stimulus Generation and Editing

We created a set of five-second audiovisual clips following standardized guidelines to maximize clarity and consistency across the archive. Each clip featured a trigger action presented continuously throughout its duration to ensure stable exposure and minimize variability in onset timing. However, for certain actions (e.g., swallowing), continuous presentation was not feasible, since these actions occur as discrete events rather than sustained behaviors. All stimuli included a clearly identifiable auditory component and a corresponding visual component, with the sound and action tightly synchronized. To reduce potential confounds, we minimized distractions by controlling background noise, omitting full human faces, and avoiding cluttered environments or extraneous features, while preserving the naturalistic quality of the stimuli. We began with the most common triggers and subsequently expanded the set to include participant-specific triggers reported by individuals with misophonia.

All recordings were captured with handheld devices in landscape orientation to ensure consistency across clips and compatibility with common display formats. Post-processing followed a standardized pipeline using ezgif.com: (1) cropping to a 4:3 aspect ratio, (2) resizing to 640 × 480 pixels to balance visual clarity with manageable file size, (3) trimming to a fixed duration of 5 seconds, and (4) converting to .mp4 format for broad compatibility across research platforms. Audio was further standardized through mono conversion, noise reduction, and loudness normalization to −23 LUFS. Each file was systematically named according to the convention *norm_mono_ResearcherID_actionMaterialNumber* to maintain traceability and facilitate integration into experimental workflows.

To ensure quality and consistency, all clips underwent review by multiple members of the research team. Reviewers verified that each stimulus contained a clearly identifiable trigger, was free from visual and auditory distractions, and accurately reflected its intended action category. This validation step complemented the standardized recording and post-processing procedures, reinforcing both clarity and comparability across stimuli. Together, these measures were designed to maximize ecological validity while maintaining experimental control, resulting in a stimulus set that is representative of real-world misophonia triggers yet optimized for use in controlled laboratory paradigms.

## Data Record

### Misophonia Audiovisual Trigger Archive (MATA)

Audiovisual stimuli were organized into 12 folders corresponding to broad categories (Figure 1): body-part sounds, mouth sounds (eating), mouth sounds (not eating), nasal-throat sounds, repetitive and continuous sounds (animal), repetitive and continuous sounds (human), repetitive and continuous sounds (non-human), rubbing sounds, rustling and tearing sounds, speech and vocalization sounds, talking sounds, and walking sounds. The distribution of files across these categories is shown in Figure 2. All audiovisual stimuli are stored in .*mp4* format. The archive is version-controlled on GitHub (https://github.com/Svetlana-Shinkareva/MATA) and its latest release is hosted on OSF https://osf.io/e7qbm/. OSF also provides a DOI for provenance tracking (10.17605/OSF.IO/E7QBM).

**Figure 1.**
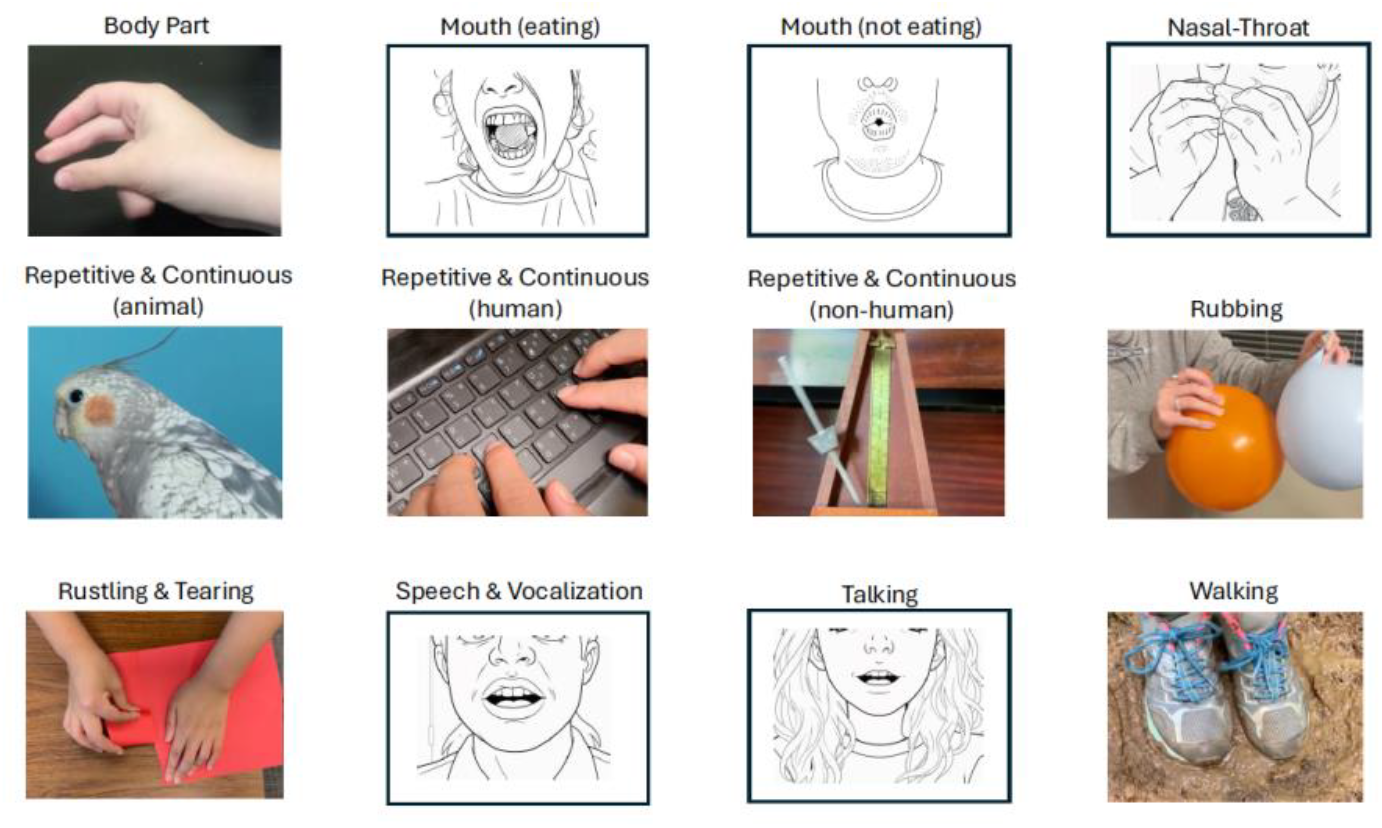
Illustration of the twelve categories of audiovisual stimuli contained in the Misophonia Audiovisual Trigger Archive (MATA), with representative still frames shown for each category. This taxonomy provides a structured framework for organizing and selecting stimuli in misophonia research. For the purposes of this preprint, human faces have been replaced with line drawings.

**Figure 2.**
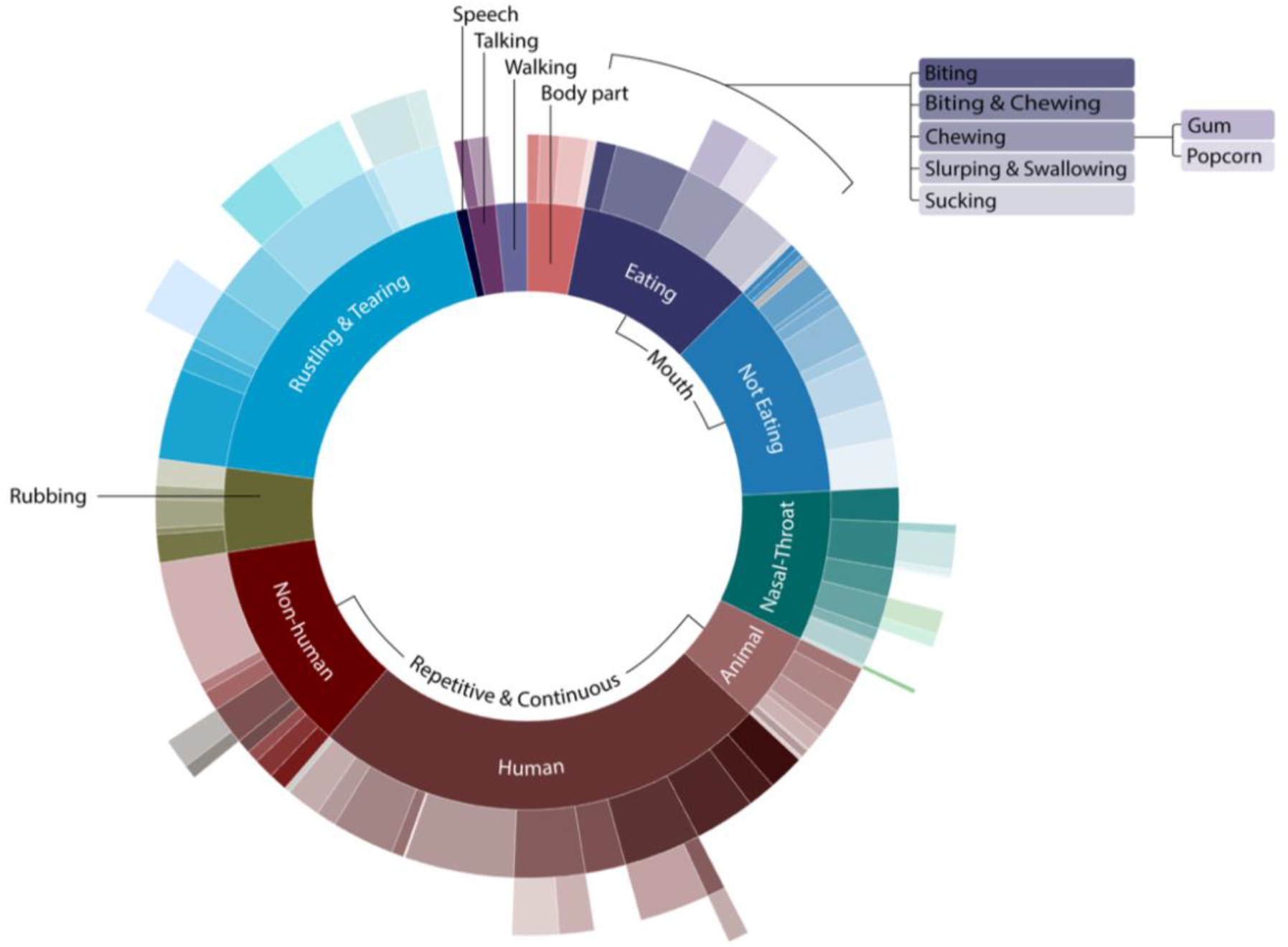
Sunburst visualization of the stimulus archive, displaying twelve primary categories (inner circle) and their hierarchical subcategories (three outer rings). This structure provides an organized overview of 1,300 stimuli in the archive and facilitates systematic selection of individualized triggers for experimental use.

### Validation data

Ratings of trigger, aversive, and non-aversive audio and visual trials from paired participants with and without misophonia are available in the MATA GitHub repository. The dataset is provided in long format as a comma-separated values file (.csv) and includes demographic information for each participant.

## Technical Validation

The goal of the technical validation was to demonstrate that the audiovisual archive reliably elicits misophonia-specific responses. We aimed to confirm that participants with misophonia show stronger distress to individually selected trigger stimuli compared to aversive stimuli across auditory and visual modalities, while control participants do not differentiate between these stimulus types.

### Participants and experimental design

The study included 26 individuals with misophonia (*M*_*age*_ = 25.18, *SD*_*age*_ = 7.14; 1 left-handed; 5 males) and 26 control participants without misophonia, matched to the misophonia group on age, biological sex, handedness, and stimulus presentation (*M*_*age*_ = 25.53, *SD*_*age*_ = 6.98; 1 left-handed; 5 males). All participants singed the informed consent approved by the institutional review board of the University of South Carolina and completed the Selective Sound Sensitivity Syndrome Scale misophonia questionnaire [S-Five; 15] to assess the presence and severity of misophonia.

During the experiment, participants were exposed to auditory-only and visual-only presentations of trigger stimuli, as well as aversive and non-aversive stimuli. A mental imagery condition was also included but is not discussed here. Participants in the misophonia group (*M*_S-Five_= 120.65, *SD*_*S-Five*_ = 37.25) rated each stimulus for subjective distress on a 4-point scale (1 = not at all, 4 = extremely). Control participants (*M*_S-Five_= 42.54, *SD*_*S-Five*_ = 10.90), for whom distress ratings were not applicable, instead rated each stimulus on perceived antisociality using the same 4-point scale. Trigger stimuli were individually selected for each participant with misophonia based on their self-reported trigger list. The total number of trigger stimuli presented, 36, was held constant across participants. However, because sufficient numbers of unique triggers with visual components were not always available, some participants were presented with repeated instances of the same stimulus. In total, 311 unique trigger stimuli were selected. To prevent confounding, aversive and non-aversive control stimuli were curated to exclude potential triggers. Each control participant was presented with the same set of stimuli as their matched misophonia participant to ensure comparability across groups.

### Results

For auditory components, participants with misophonia rated trigger stimuli as significantly more distressing than aversive stimuli, *t*(25) = 11.92, *p* < .001, 95% CI [1.03, 1.59], *d* = 2.25. A similar pattern was observed for the visual components, with trigger stimuli rated as significantly more distressing than aversive stimuli, *t*(25) = 8.82, *p* < .001, 95% CI [0.84, 1.53], *d* = 1.67. In contrast, control participants rated trigger and aversive stimuli as comparably antisocial for both auditory, *t*(25) = 0.03, *p* = 1.000, 95% CI [-0.20, 0.27], and visual components, *t*(25) = 1.29, *p* = 0.622, 95% CI [-0.32, 0.11] (Figure 3). These results demonstrate that participants with misophonia reliably rated trigger stimuli as more distressing than aversive stimuli across both auditory and visual conditions, confirming that the trigger stimuli evoke misophonia-specific responses.

**Figure 3.**
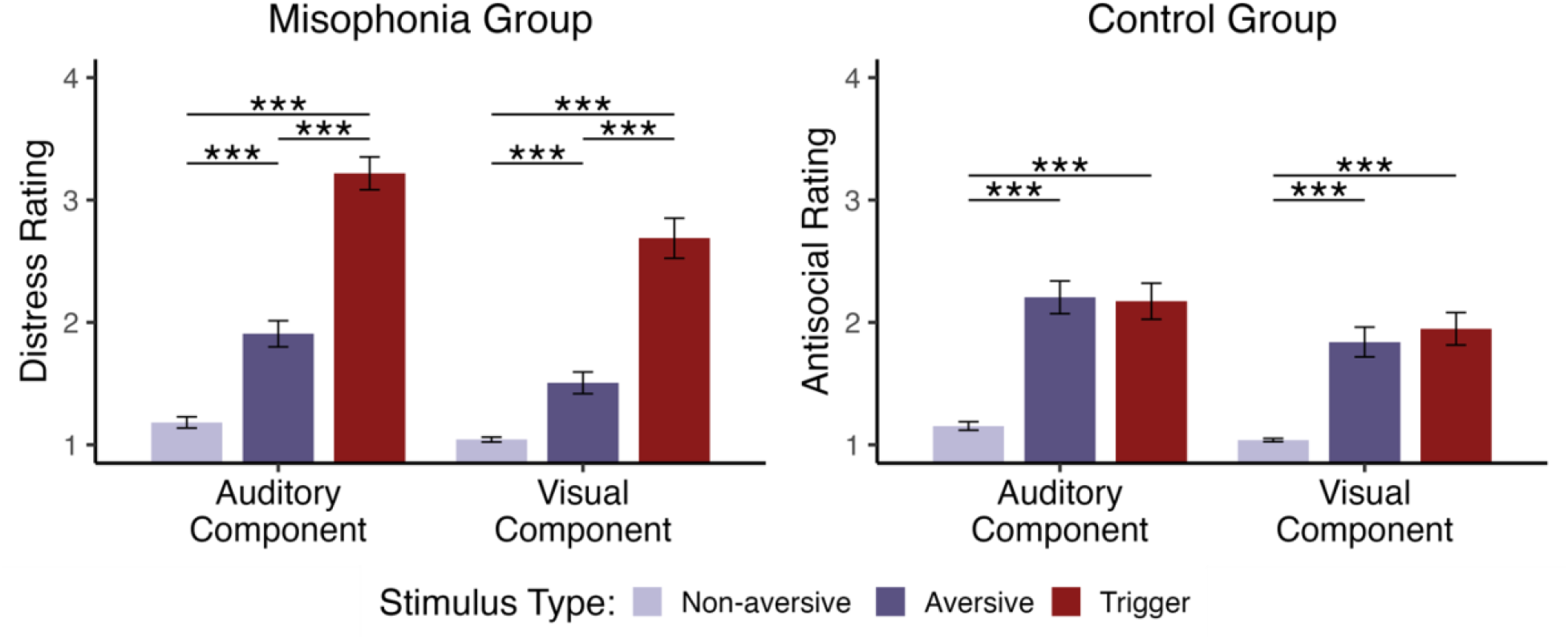
Mean (±SE) subjective distress ratings for the misophonia group and perceived antisociality ratings for the control group, shown separately for auditory and visual components of non-aversive, aversive, and trigger stimuli. Participants with misophonia consistently responded more strongly to trigger than to aversive stimuli across auditory and visual conditions, whereas control participants did not differentiate between the two. ^***^ p < .001

## Acknowledgement

This project was funded by the Misophonia Research Fund. We thank our research team for their contributions to stimulus generation and for the valuable discussions that informed the development of this work.

## Author contributions

S.V.S. and S.O. designed and supervised the study. S.O., K.P., D.S., and A.P. organized and edited the stimuli. S.V.S., S.O., K.P., and C.O. drafted and revised the manuscript. All authors discussed the results, contributed to the writing, reviewed the manuscript, approved the final version, and agreed to be accountable for all aspects of the work.

